# Benchmarking Genomic Variant Calling Tools in Inbred Mouse Strains: Recommendations and Considerations

**DOI:** 10.1101/2025.05.28.656711

**Authors:** Alexis Garretson, Laura Blanco-Berdugo, Aleisha Roberts, Beth L. Dumont

## Abstract

With the growing affordability of whole genome sequencing, variant identification has become an increasingly common task, but there are many challenges due to both technical and biological factors. In recent years, the number of software packages available for variant calling has rapidly increased. Understanding the benefits and drawbacks of different tools is important in setting leading practices and highlighting limitations. These considerations are crucial in model organism research, as many variant calling programs assume outbred genomes and implicit heterozygosity, which may not apply to inbred laboratory models. Here, we present an analysis of variant calling tools and their performance in the simulated genomes of the C57BL/6J inbred laboratory mouse and nine non-reference laboratory strains. Our findings reveal a tradeoff between the recall and precision of tools. Balancing these considerations, we show that an optimal call set is obtained by using an ensemble approach, but specific variant calling recommendations vary by strain and analytical goals. Further, we highlight filters improving the performance of different variant calling tools, both for the discovery of rare variants and in the discovery of strain polymorphisms. In summary, our work provides best practices for calling and filtering genomic variants in inbred organisms, particularly laboratory mice.

**Article Summary:** Identifying mutations and rare genetic variants is a central task for modern genomics. Many computational tools exist for variant detection, but their performance varies across diverse applications. Further, few variant calling tools have been benchmarked against inbred genomes, which are commonly used for research. To address this, we evaluated five variant calling tools using simulated data from ten diverse inbred mouse strains. We show that variant detection, recall, and precision vary across tools and mouse strains, and that an ensemble approach improves confidence in detected mutations. Our findings offer a set of best practices for variant calling in inbred organisms across diverse analytical applications.

## Introduction

Despite their crucial importance for clinical research and basic biology, accurately identifying and calling single-nucleotide polymorphisms (SNPs) from high-throughput sequencing data remains an imperfectly solved challenge. Sequencing errors, alignment artifacts, biological variation, and noise can all give rise to false positive or false negative SNP calls (Li 2014; Krusche et al. 2019; Stoler and Nekrutenko 2021; Wang et al. 2023). These errors can have significant impacts on the interpretation and accuracy of research findings, as SNPs are critical inputs to GWAS and other downstream analyses. SNPs also have significant functional importance. SNPs have been causally linked to numerous diseases and phenotypic traits (Frazer et al. 2007; Wray et al. 2007; Altshuler et al. 2010; MacArthur et al. 2017) and can modify the efficacy of different therapeutic treatments in different individuals (Mackay et al. 2009; Alexandrov et al. 2013; Goddard et al. 2016; Lewis and Vassos 2020). The genomic distribution of SNPs also holds critical information about genome evolutionary history, including the history of selective sweeps and demographic events (Galtier et al. 2000; Schork et al. 2000; Emerson et al. 2001; Shakya et al. 2020).

Numerous software packages and bioinformatic pipelines have been developed for SNP calling from whole genome sequences (DePristo et al. 2011; DePristo et al. 2011; Li 2011; Garrison and Marth 2012). These approaches have been at the forefront of scientific research in recent years, with availability and usage of tools varying over time (Figure 1). Different variant calling tools employ diverse algorithms, statistical approaches, and filtering strategies (Table 1). These fundamental distinctions mean that different tools can produce different variant call sets from the same input sequencing data. While this point is tacitly recognized, limited guidance exists in the scientific literature to aid investigators in selecting the optimal variant calling tool or combination of tools in view of diverse analytical applications.

**Figure 1.**
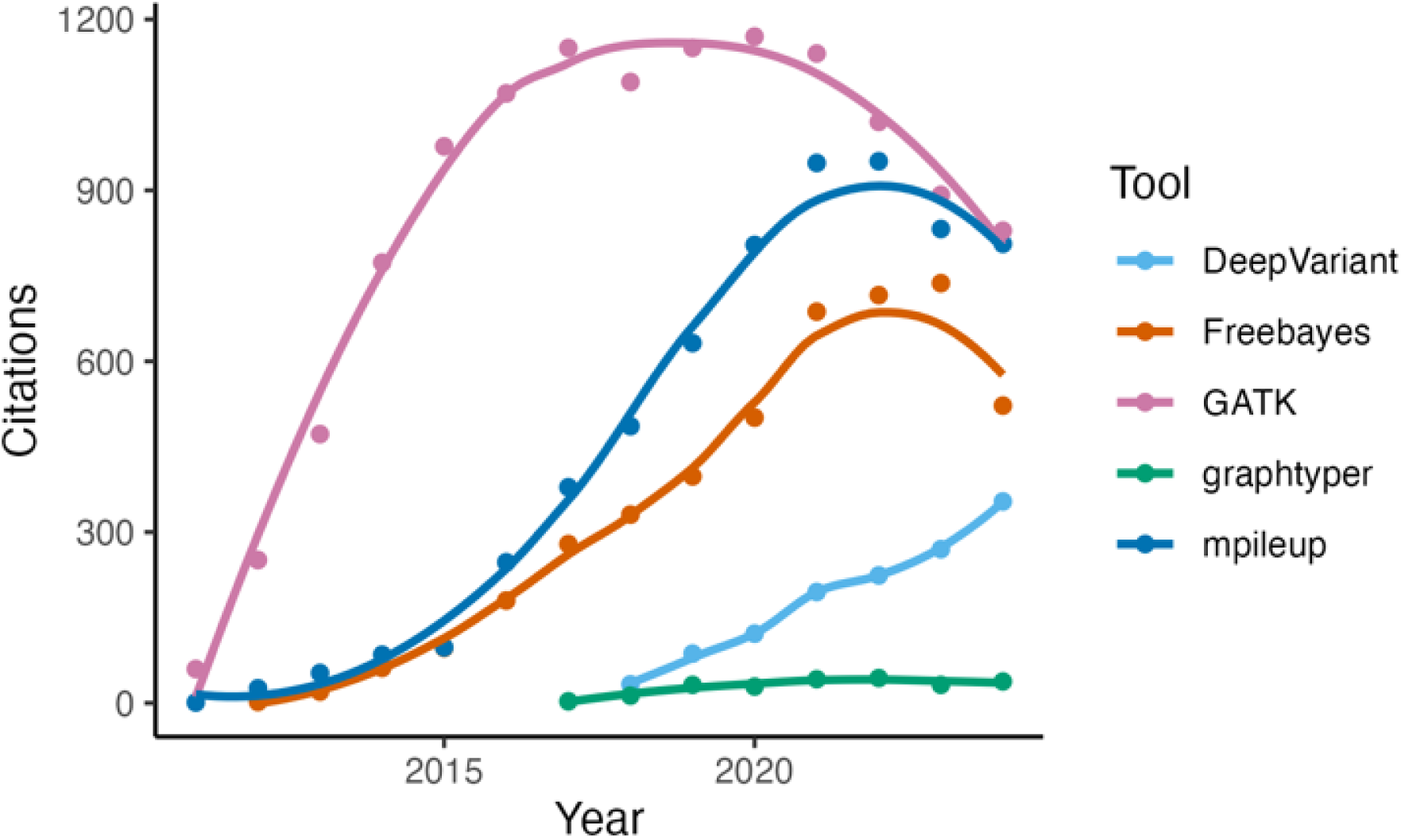
Counts of the number of papers citing each of the four variant calling tools evaluated in this work over a 12-year span from 2011 - 2024 (source: Google Scholar 25-FEB-2025).

**Table 1.** Variant calling tools evaluated in this study with software version and algorithm approach.

|  |  |  |
| --- | --- | --- |
| Variant tool | Version | Algorithm |
| Freebayes (Garrison and Marth 2012) | v1.3.1 | Haplotype-based |
| GATK (DePristo et al. 2011) | v4.2.3.0 | Haplotype-based |
| bcftools /mpileup (Li 2011) | v1.9 | Site align-based |
| DeepVariant (Poplin et al. 2018) | v1.2.0 | Deep neural network |
| graph typer (Eggertsson et al. 2019) | v2.7.7 | Graph-based |

Consequently, researchers are often faced with uncertainty when selecting the most appropriate tool, with limited knowledge of the comparative strengths and weaknesses of competing approaches. Existing benchmark comparisons of variant calling tools use actual or simulated data from human genomes or other outbred organisms (O’Rawe et al. 2013; Pirooznia et al. 2014; Bian et al. 2018; Yao et al. 2020; Lefouili and Nam 2022). Additionally, the statistical models employed by many variant calling tools are developed and benchmarked against outbred samples, and therefore rest on assumptions about levels of heterozygosity in diploid genomes (Pirooznia et al. 2014; Warden et al. 2014; Krusche et al. 2019; Barbitoff et al. 2022). Thus, current best practice workflows for detecting polymorphic SNPs may not be generalizable to researchers working in highly inbred or haploid model systems.

In this paper, we present a comprehensive analysis of five widely used variant calling tools (DeepVariant, GATK, Graphtyper, Mpileup, Freebayes) and their performance in simulated genomes of the reference inbred laboratory mouse strain C57BL/6J (B6) and in nine non-reference inbred strains: 129S1/SvImJ (129), A/J (AJ), CAST/EiJ (CAST), NOD/ShiLtJ (NOD), NZO/HiltJ (NZO), PWK/PhJ (PWK), WSB/EiJ (WSB), SPRET/EiJ (SPRET), and DBA/2J (DBA). We evaluate the accuracy and sensitivity of different variant calling approaches and provide benchmark and practical guidance for variant calling using inbred genomes.

## Materials and Methods

### Mutation and read generation

Mutations were simulated on the mouse reference genome (GRCm39) using simuG (v1.0.0; Yue and Liti 2019). Ten independent replicates of chromosome 1 were generated, each containing 5,000 randomly placed fixed differences at a transition/transversion ratio of 1.5 compared to the reference genome (Figure 2). simuG produced two key outputs: (1) a text file recording the genomic positions of the simulated SNPs and (2) a FASTA file in which these SNPs were inserted at their respective locations.

**Figure 2.**
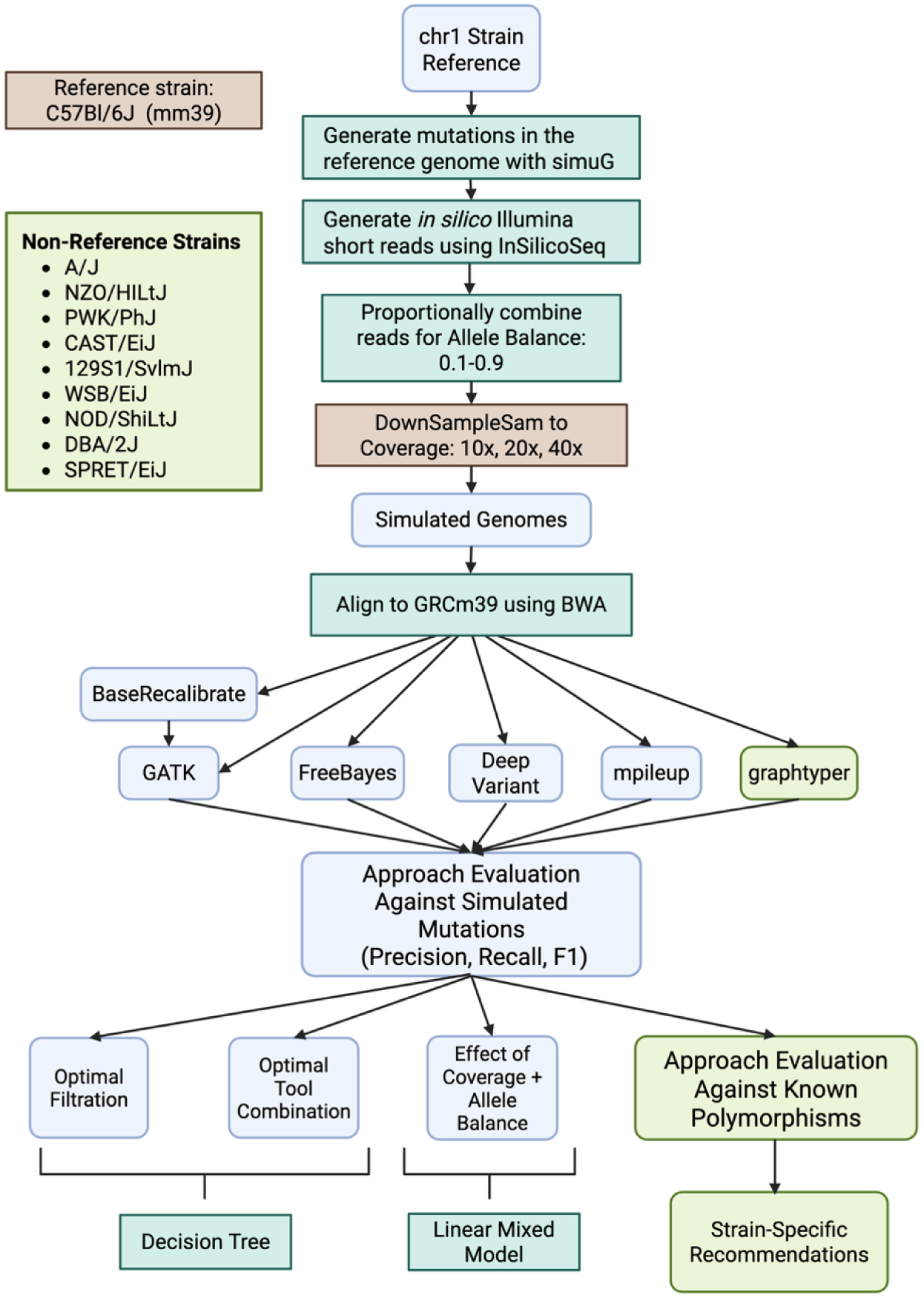
Schematic for experimental design. Boxes in light green indicate steps only performed for the non-B6 strains while the boxes in orange indicate steps done for just B6. All other portions of the protocol were repeated for both sample sets.

Next, InSilicoSeq (v1.5.3; Gourlé et al. 2019) was used to generate simulated Illumina paired-end 150 bp sequencing reads, applying a NovaSeq error model. Sequencing was simulated at a uniform coverage of 40× by using the Lander/Waterman equation (Lander and Waterman 1988)

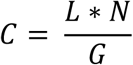

Where C is the intended coverage, L is the read length, N is the number of reads, and G is the genome size. To simulate at coverage 40× for only chr1, we set G equal to the chr1 length in bp, leading to 52.0M reads. To evaluate the impact of allele balance (AB) on SNP calling, the output FASTA files from simuG were combined with the unmodified reference chromosome 1 FASTA. InSilicoSeq was then used to generate FASTQ files reflecting ABs ranging from 0.1 to 0.9 by proportionally mixing variant-supporting reads with reference reads. This process ensured that all resulting FASTQ files maintained a coverage of 40× across replicates and AB conditions. To further investigate the effects of coverage, the 40× coverage BAM files were randomly down-sampled to approximately 10× and 20× coverage using Picard Tools (v2.23) DownsampleSam. This tool employs a probabilistic approach to retain the desired proportion of input reads, ensuring consistent coverage reduction while preserving read distribution characteristics.

For the nine non-reference strains—WSB/EiJ, 129S1/SvImJ, A/J, CAST/EiJ, DBA/2J, NOD/ShiLtJ, NZO/HlLtJ, PWK/PhJ, and SPRET/EiJ—mutations were simulated on each strain-specific assembly (Lilue et al. 2018, for accession numbers see Table S1). For each non-reference strain genome, we generated ten replicates of 5,000 variants introduced into each chr1 sequence at 40×. Despite slight differences in the chr1 genome length for the non-reference strains, we simulated the same number of reads as we did for B6, meaning that coverage values fluctuate slightly across strains. To determine the corresponding location on the GRCm39 reference, coordinate conversion was performed via Picard *LiftOverVcf* (v2.23) with chain files generated by the UCSC Genome Browser (Table S1).

### Variant Calling Tool Application

We aligned all simulated sequencing reads to the GRCm39 reference genome using *BWA* (0.7.17) (Li and Durbin 2009) and performed variant calling. Four variant calling tools (DeepVariant, FreeBayes, GATK, and Mpileup) were applied to B6 simulated data, and an additional tool, Graphtyper, was used for variant calling in the nine non-reference strains. For all variant calling tools except graphtyper, variants were called individually for each set of simulated reads. For graphtyper, calling was performed jointly across all strains for each replicate (for example, jointly on all nine strains for simulation replicate produced under AB 0.5 and 40× coverage). This distinction owes to the fact that a pangenome approach is most beneficial for providing a comprehensive representation of genomic diversity across a population, rather than just capturing the genetic content of a single individual or strain (Eggertsson et al. 2017; Sibbesen et al. 2018).

Mpileup (executed under the command “bcftools call” in bcftools v 1.9-1) and Freebayes (v 1.3.1) were run with default parameters. SNPs were extracted and subjected to the following post-hoc filters using vcflib (v1.0.10): "QUAL > 1 & QUAL / AO > 10 & SAF > 0 & SAR > 0 & RPR > 1 & RPL > 1". The DeepVariant callset was generated using the WGS model in DeepVariant (v 1.2.0) and limiting calling to the simulated region with the flag “–regions chr1”. GATK (v 4.2.2.0) variant calling proceeded in multiple steps. First, all read alignments to the GRCm39 reference were regrouped, and duplicated reads were marked using gatk MarkDuplicatesSpark. Resulting bams were recalibrated using gatk BaseRecalibrator and gatk ApplyBQSR. Next, SNPs were called using gatk HaplotypeCaller, limiting the call region to chr1 (-L chr1). We applied GATK-recommended hard filters: (“QD < 2.0 || FS > 20.0 || MQ < 40.0 || SOR > 3.0 || MQRankSum < -2.0 || MQRankSum > 4.0 || ReadPosRankSum < -3.0 || ReadPosRankSum > 3.0. Graphtyper variant calling was performed using default parameters but limited to just the chr1 region (--region=chr1). Output variants were filtered to just the PASS variants using default filters of ABHet < 17.5%, ABHom < 90%, QD < 6, QUAL score < 10, or AAScore < 0.5. Variants with a reference call (e.g., 0/0), or where the alternative call overlapped a deletion (“ALT=*”) were also removed, and any remaining multiallelic sites were split to biallelic records using bcftools norm -m - both.

### Evaluation of Resulting Call Sets

To evaluate the performance of each variant calling tool on our B6 simulation-based callsets, we annotated each SNP call as a false positive (i.e., called but absent from the simulated variants), false negative (i.e., present in the simulated data but not called by the tool), or true positive (i.e., called and present in the simulated data). Using the number of variants of each type, we calculated the precision, recall, and F1 score for each variant calling tool, for each replicate, and for each AB and coverage. Significance between tools and simulation parameters was assessed using Kruskall-Wallis tests.

Pair-wise performance of tools across metrics was assessed by Wilcoxon signed rank tests. The relationship between sequencing metrics and variant properties was evaluated using a linear regression model. Because the relationship between AB and each variant calling performance metric appeared logarithmic, we applied a natural logarithm transformation to the AB variable. While we initially fit a mixed-effects model with replicate as a random effect, the variance estimates for the replicate-level random effect were near zero, indicating minimal variation attributable to this factor. Additionally, the model fit was singular, indicating overparameterization. As a result, we simplified the model by omitting the random replicate effect, leading to the final model:

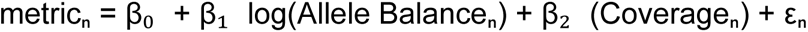

where metricₙ represents either precision or recall for observation *n*, β₀ is the intercept, β₁ and β₂ are regression coefficients for the log-transformed AB and coverage, respectively, and εₙ represents the residual error. The model was fit using the lm() function in R (v4.4.0).

For the diverse strains, we consider only a single, fixed coverage value for each strain, and independently performed modeling fitting for each strain. As with the reference-strain model, we observed that the replicate variance was approximately 0 across all strains leading to singular model fits, justifying exclusion of replicate as a random factor in the final model:

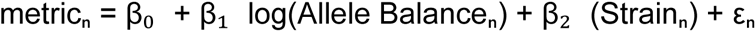

where metricₙ represents either precision or recall for observation *n*, β₀ is the intercept, β₁ is the regression coefficient for the log-transformed AB, β_2_ is the genetic distance from the reference measured as the total number of fixed SNP differences on chr1 between the focal strain and the GRCm39 reference, and εₙ represents the residual error. Model fitting was performed using the lm() function in R.

### Machine Learning of Optimal Tool Combination and Filtration Strategies

We used rpart (Therneau et al. 2023) via the tidymodels (Kuhn and Wickham 2023) package to generate a classification decision tree to determine the best combination of variant calling tools and to assess whether additional filtering could improve SNP calling performance. For this purpose, we designated 25% of all called variants as a testing set and the remaining 75% as a training set. We generated CART (<u>c</u>lassification <u>a</u>nd <u>r</u>egression tree) decision trees based on fitting a model predicting the probability a variant was in the simulation dataset as a function of whether a given variant calling tool had correctly called that site.

To evaluate the performance of variant calling in the nine non-B6 strains, we followed the above procedure, with some modifications to include tool performance in recovering known strain-specific polymorphisms (Keane et al 2011). We built CART classification trees for each tool, AB, and strain combination using the following features: (1) the genotype of the called variant, (2) whether the site was included in the set of known SNPs for that strain, (3) the output quality score for the variant, (4) the RepeatMasker family-level annotation (based on the GRCm39 annotations), and (5) whether the variant fell into a structurally variable region of that strain’s genome (as annotated by Ferraj et al. 2022 for all strains excluding SPRET) and, if so, SV type (insertion or deletion) and the SV length, and (6) whether the variant fell into a set of genes known to be rapidly evolving and highly copy number variable in mice (vomeronasal receptors, olfactory genes, and zinc-finger proteins) (Niimura et al. 2024).

## Results

### Precision and Recall Trade-offs in Variant Calling Performance

We began by assessing the performance of four variant calling tools using simulated sequence reads derived from the mouse reference genome. This scenario represents a “best case” as sequence divergence from the reference is restricted to sites of simulation-induced point mutations, minimizing possible variant calling artifacts due to read mis-mapping. Recall, precision, and F1 scores varied significantly across variant calling tools and simulation conditions (Figure 3, Table S2). Under the set of simulation conditions where variant calling is expected to be the most accurate (i.e., AB = 0.9 and coverage = 40x), we observed negligible differences in precision performance between tools. Under these ideal conditions, DeepVariant was the only caller with significantly lower precision than other callers, but the magnitude of this difference was slight (precision = 0.999±0.0003). However, at lower coverage values, DeepVariant’s reduced precision was counter-balanced by higher recall (0.995 ± 0.001; Mpileup = 0.993 ± 0.001; Freebayes = 0.977±0.001; GATK = 0.901±0.003).

**Figure 3.**
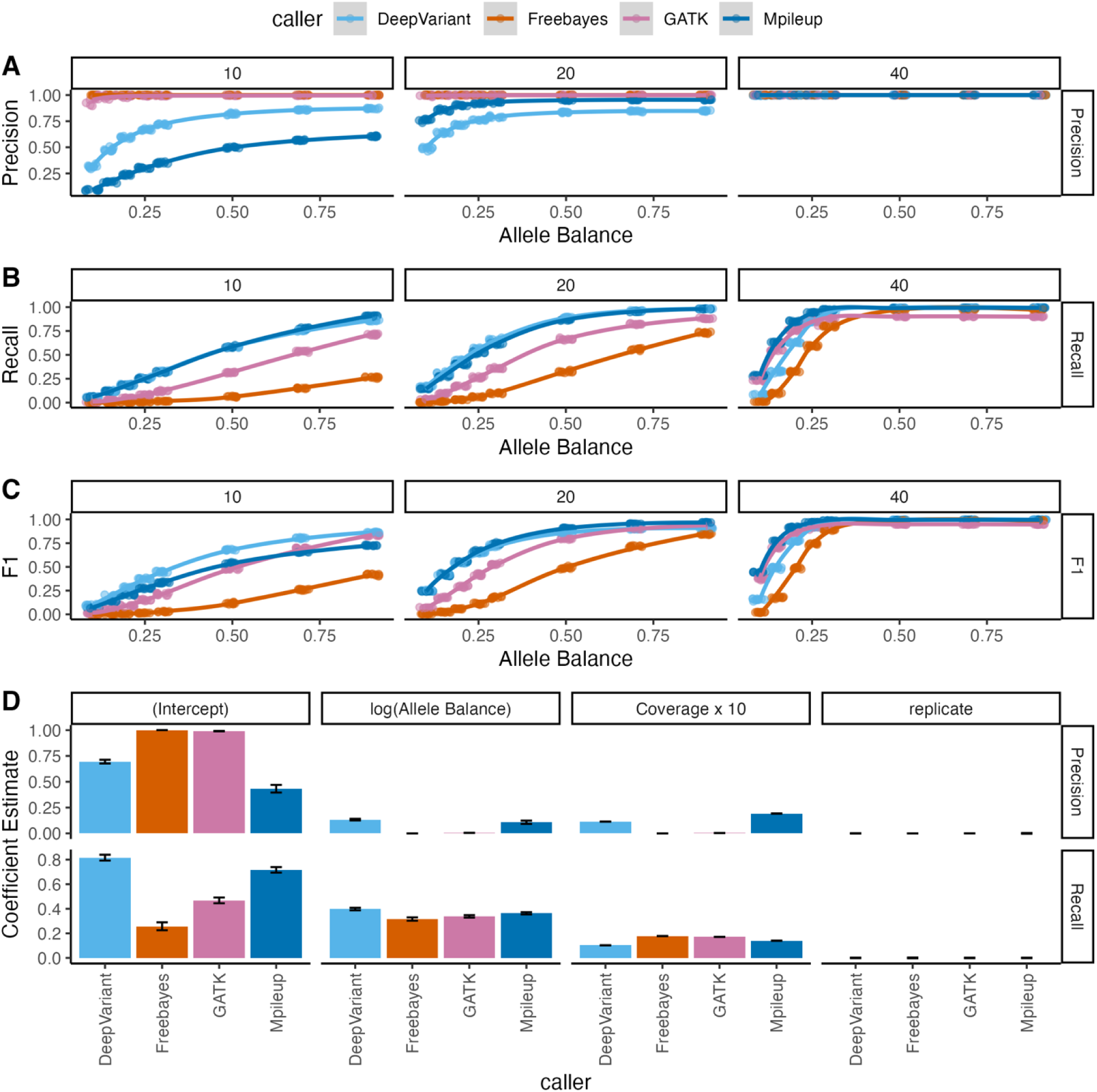
Variant calling tool performance comparison using simulated data based on the B6 reference genome. Tools were evaluated across eight allele balance conditions and 3 sequence depth values using (A) precision, (B) recall, and (C) F1 score. Comparison of coefficient estimates from linear models assessing the effect of log-transformed allele balance and coverage on precision and recall for the detection of simulated variants using different variant calling tools. For ease of comparison, the sequence depth values are represented as increments of 10, meaning that coefficient estimates correspond to the increase in the target metric with a 10x increase in coverage.

Overall, the performance across replicates for each simulation condition was highly consistent (average variance: recall=2.9×10^-5^, precision=3.4×10^-5^, and F1 = 2.9×10^-5^, Figure 3A-D), indicating that the specific locations of the simulated mutations had negligible effect on the general performance of each variant calling tool. However, not unexpectedly, our results indicate that coverage and AB substantially impacted the performance of each evaluated tool (Figure 3B, C). While increased AB and higher coverage generally improved performance, the magnitude of the improvements in recall and precision varied by caller (Table 2, Figure 3D).

**Table 2.** Results of a tool-specific linear regression to evaluate performance metric (precision or recall) as a function of both the natural log of allele balance and coverage. Coefficients and the intercept are reported ± 1 standard error, with t and P values. Additionally, the R2, adjusted for two predictors, and the model P value are reported.

| Caller | Name | Intercept | log(Alelle Balance) | Coverage | R<sup>2</sup><sub>adj</sub> | P Value |
| --- | --- | --- | --- | --- | --- | --- |
| DeepVariant | Precision | 0.695±0.015<br>t=47.16, P=5×10<sup>-121</sup> | 0.133±0.008<br>t=17.43, P=6×10<sup>-43</sup> | 0.011±0<br>t=26.04, P=4×10<sup>-70</sup> | 0.80 | 5×10<sup>-85</sup> |
| | Recall | 0.398±0.01<br>t=39.51, P=7×10<sup>-105</sup> | 0.398±0.01<br>t=39.51, P=7×10<sup>-105</sup> | 0.01±0.001<br>t=17.89, P=2×10<sup>-44</sup> | 0.89 | 2×10<sup>-113</sup> |
| Freebayes | Precision | 1±5×10<sup>-16</sup><br>t=2E+15, P=0 | 5×10<sup>-16</sup>±3×10<sup>-16</sup><br>t=1.6, P=1 | 2×10<sup>-17</sup>±2×10<sup>-17</sup><br>t=1.11, P=1 | 0.50 | 3×10<sup>-36</sup> |
| | Recall | 0.257±0.026<br>t=9.78, P=8×10<sup>-18</sup> | 0.317±0.014<br>t=23.2, P=2×10<sup>-61</sup> | 0.018±0.001<br>t=22.82, P=2×10<sup>-60</sup> | 0.82 | 4×10<sup>-88</sup> |
| GATK | Precision | 0.993±0.002<br>t=525.93, P=0 | 0.005±0.001<br>t=5.43, P=3×10<sup>-6</sup> | 4×10<sup>-4</sup>±6×10<sup>-5</sup><br>t=6.56, P=8×10<sup>-9</sup> | 0.23 | 2×10<sup>-14</sup> |
| | Recall | 0.467±0.019<br>t=24.6, P=8×10<sup>-66</sup> | 0.338±0.01<br>t=34.37, P=1×10<sup>-92</sup> | 0.017±0.001<br>t=30.46, P=2×10<sup>-82</sup> | 0.9 | 1×10<sup>-118</sup> |
| Mpileup | Precision | 0.433±0.03<br>t=14.25, P=3×10<sup>-32</sup> | 0.108±0.016<br>t=6.85, P=2×10<sup>-9</sup> | 0.019±0.001<br>t=21.32, P=1×10<sup>-55</sup> | 0.68 | 3×10<sup>-59</sup> |
| | Recall | 0.716±0.018<br>t=39.48, P=8×10<sup>-105</sup> | 0.363±0.009<br>t=38.65, P=6×10<sup>-103</sup> | 0.014±0.001<br>t=26.09, P=3×10<sup>-70</sup> | 0.9 | 4×10<sup>-120</sup> |

### Ensemble Approaches Improve Variant Calling Performance, Particularly at Low Coverage

To better understand the performance differences across variant calling tools, we next evaluated the intersections of variant call sets across tools at each coverage (10x, 20x, 40x) and at 3 representative ABs (0.1, 0.5, and 0.9; Figure 4A, Figure S1, S2). We uncover nuanced patterns of agreement and divergence between callers that are not captured by overall precision and recall estimates. For example, even at high coverage, the vast majority of variants simulated at AB 0.1 were missed by all callers (Figure 4A). While performance improves as AB increases, a large share of simulated variants were still missed by all tools, especially at low coverage (Figure 4A, Figure S1, S2). On the other hand, the majority of false positive calls are unique to a single tool, rather than shared across callers (Figure 4A). Of those false positive variants that are detected by multiple callers, most are shared by Mpileup and DeepVariant, suggesting common error profiles associated with these two tools (Figure 4A). GATK recovered few unique true positives, but a relative excess of these unique true variant calls was recovered under low AB conditions compared to other callers. GATK may have unique performance gains for detecting rare, low AB variants, particularly in combination with other callers that recover a greater share of variants under these conditions (Figure 4A, Figure S1, S2).

**Figure 4.**
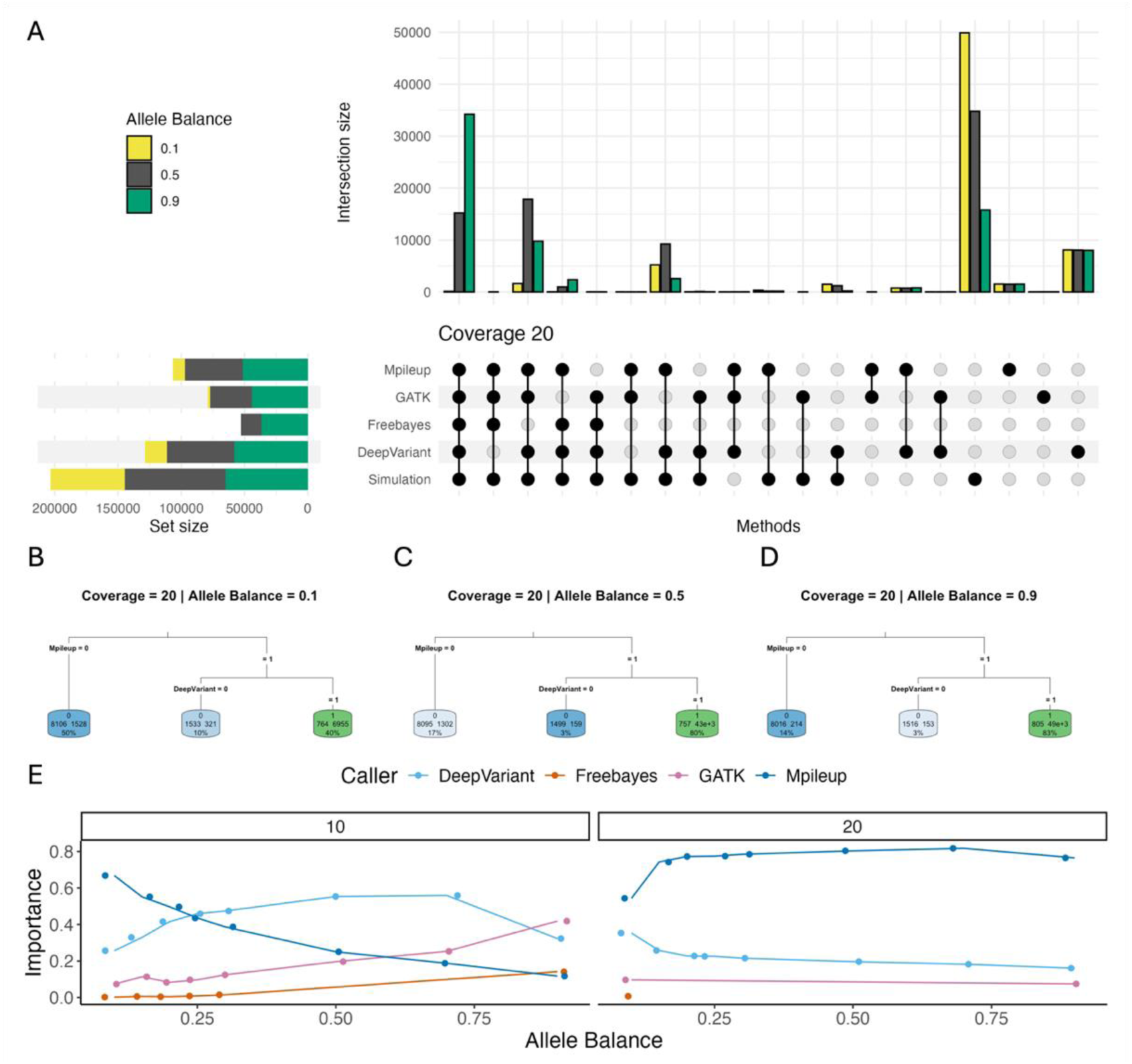
Comparisons of SNP call sets across tools. (A) Upset plot of variant call sets at 20x coverage and allele balances 0.1, 0.5, and 0.9. Bar graphs show the intersection sizes between each combination of tools, with each colored bar showing the size of that set at the given allele balance. Set sizes to the left of the upset chart panel indicate the number of variants recovered by that tool under the simulated allele balance condition. (B,C,D) Classification tree predicting variant status based on the optimal combination of variant calling tools under different allele balance conditions. Simulated variants are represented as “1”, and false positives are indicated by “0”. Splitting criteria are displayed on the connecting branches, indicating the decision rule applied to divide the data. Terminal (leaf) nodes show the final predicted value and sample size for each group of observations. Node color intensity represents the relative probability that an observation in that node belongs to the predicted class, with green indicating a higher probability (closer to 1) and blue indicating a lower probability (closer to 0). Darker colors correspond to more confident predictions, while lighter colors indicate greater uncertainty or class mixing within the node. (E) Importance of variant calling tools in classification trees across different allele balance and coverage levels. Each line connects points representing the importance of a specific variant calling tool (indicated by line color) in classification trees predicting the true/false positive status of the variant. Panels are faceted by simulated sequencing coverage.

We next used a decision tree framework to find the optimal ordering and combination of variant calling tools to apply in pursuit of a high-quality variant call set (Figure 4B-E). To make this discrimination, we relied on the importance values, which quantify how much each feature (in this case, each variant calling tool) contributes to correctly discriminating between true and false positives, with higher importance values indicating greater influence. At 10x coverage, importance values varied across simulated AB values, with no single tool consistently dominating across all conditions (Figure 4E). Mpileup had high importance at lower ABs but importance declined with increasing AB, whereas DeepVariant had intermediate importance at high and low AB but higher importance at intermediate AB (Figure 4E). GATK had relatively low importance at low AB, but importance rose steadily with increasing AB (Figure 4E). In contrast, at 20x coverage, importance scores stabilized across AB, with Mpileup having the highest importance across all conditions, followed by DeepVariant. GATK and Freebayes were only retained in a small number of models with extreme AB (0.1 and 0.9), and contributed little to overall model performance (Figure 4E). At a depth of 40x, all final trees had no supported splits, indicating that the most effective approach for maximizing true positive rate was to use the union of all four variant calling tools (Figure S3).

For each simulation condition, we assembled a high-quality variant call set in accordance with the optimal decision tree. Strikingly, for AB > 0.25, there is little difference in the precision of these call sets across coverage values (Figure S4). This finding suggests that the reduced performance of variant calls on low coverage datasets can be overcome, at least to some extent, by using ensemble calling approaches. In contrast, recall performance exhibits strong dependence on both coverage and AB across these optimized variant call sets, with performance gains incurred for increasing values of both simulation variables (Figure S4).

### Sample Divergence From The Reference Genome is Associated with High Numbers of False Positive SNP calls

We evaluated five variant calling tools (DeepVariant, GATK, Graphtyper, Mpileup, Freebayes) across nine genetically diverse inbred mouse strains − NZO, NOD, AJ, 129, SPRET, CAST, PWK, WSB, DBA − each with varying degrees of genetic divergence from the reference B6 strain (Figure 5A,B). Each combination of strain, AB threshold, and variant calling tool was analyzed across 10 independent simulation replicates. Because mutations were simulated against strain-specific reference genomes, coordinates were first lifted from the original strain reference to the GRCm39 reference (see Methods). The proportion of successfully lifted-over variants was high for the 8 *Mus musculus* strains, ranging from an average of 96% in PWK to 98.2% in WSB (Figure 5C), but low for the *Mus spretus* strain, SPRET (44.7%; range of 2,199 – 2,235 variants successfully lifted over across simulation replicates). Overall, we observe poor variant calling performance in SPRET, with very low recall and precision across all simulation conditions (Figure 5 D,E,F). These outcomes presumably reflect the high genetic divergence of SPRET from the GRCm39 reference, with approximately 1 sequence variant every ∼100 bp (Dejager et al. 2009).

**Figure 5.**
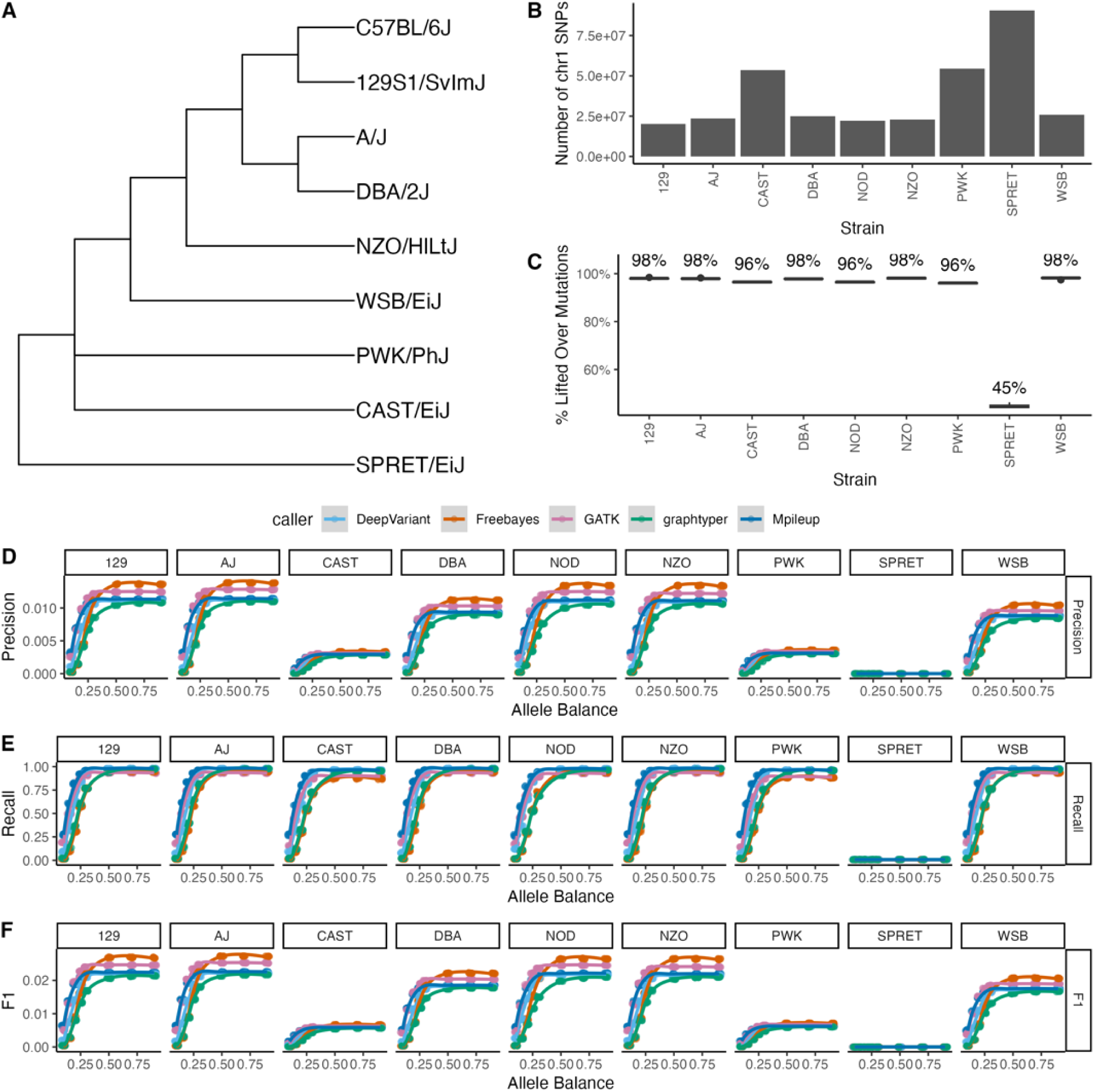
(A) Cladogram of the relationships between inbred strains, adapted from Dumont et al. 2024. (B) The number of chromosome 1 SNPs separating the surveyed strain from the GRCm39 reference. (C) The percent of the 5,000 simulated mutations in the strain-specific reference that successfully lifted-over to chromosome 1 coordinates on GRCm39. Variant calling tool performance comparison across all allele balance and strains by (D) precision, (E) recall, and (F) F1 score.

For all surveyed strains, variant call sets were dominated by false-positive calls and the resulting precision estimates were generally quite low (Figure 5D-F, Table S3). Overall, the classical laboratory strains have higher precision than CAST, PWK, and WSB (Figure 5D), a trend similarly mirrored in F1 scores (Figure 5F). However, while the precision was low, the majority of simulated mutations were detected in each call set, with recall values close to 1 at intermediate and high ABs (Figure 5E). SPRET remains an exception, with uniformly low recall across simulated ABs. For the majority of strains, Mpileup, DeepVariant, and GATK reach near perfect recall for AB >0.3, whereas graphtyper and FreeBayes do not reach similar recall performance until AB>0.5 (Figure 5E). Taken together, our findings indicate that SNP variant calling in diverse inbred strains is challenged by large numbers of false positive calls, even though most fixed variants are detected.

### Variant Calling Performance Declines as Strain Divergence from the Reference Sequence Increases

We next adopted a modeling approach to evaluate the impact of AB and strain divergence from the reference on SNP calling performance. Our results show that the intercept of precision was variable for different variant callers, with the lowest precision in graphtyper and the highest in Freebayes and GATK (Figure 6A, Table S4). (Note that, because AB is log-transformed in these models (see Methods), the intercept represents tool performance when AB is 1 and the focal strain is either identical to the reference [in models with chr1 SNP count modelled as a continuous variable]) or 129 [in models with strain treated as a factor]). However, as in the B6 results, these trends were reversed to some degree in recall performance, with DeepVariant having the highest intercept in our recall model (Figure 6A). Intriguingly, these results were not recapitulated in models evaluating caller performance at known fixed sites between strains. Instead, we report much lower recall and precision across all tools, with FreeBayes and GATK exhibiting especially poor performance (Figure 6B). AB did not contribute uniformly to increasing model performance across variant calling tools, although the relative magnitude of the performance gains for each tool were similar for both precision and recall (Figure 6A). Notably, FreeBayes and Graphtyper enjoy the largest performance gains with increasing ABs. As expected, the AB of the simulated mutations did not impact our ability to detect fixed strain-specific SNPs (Figure 6B).

**Figure 6.**
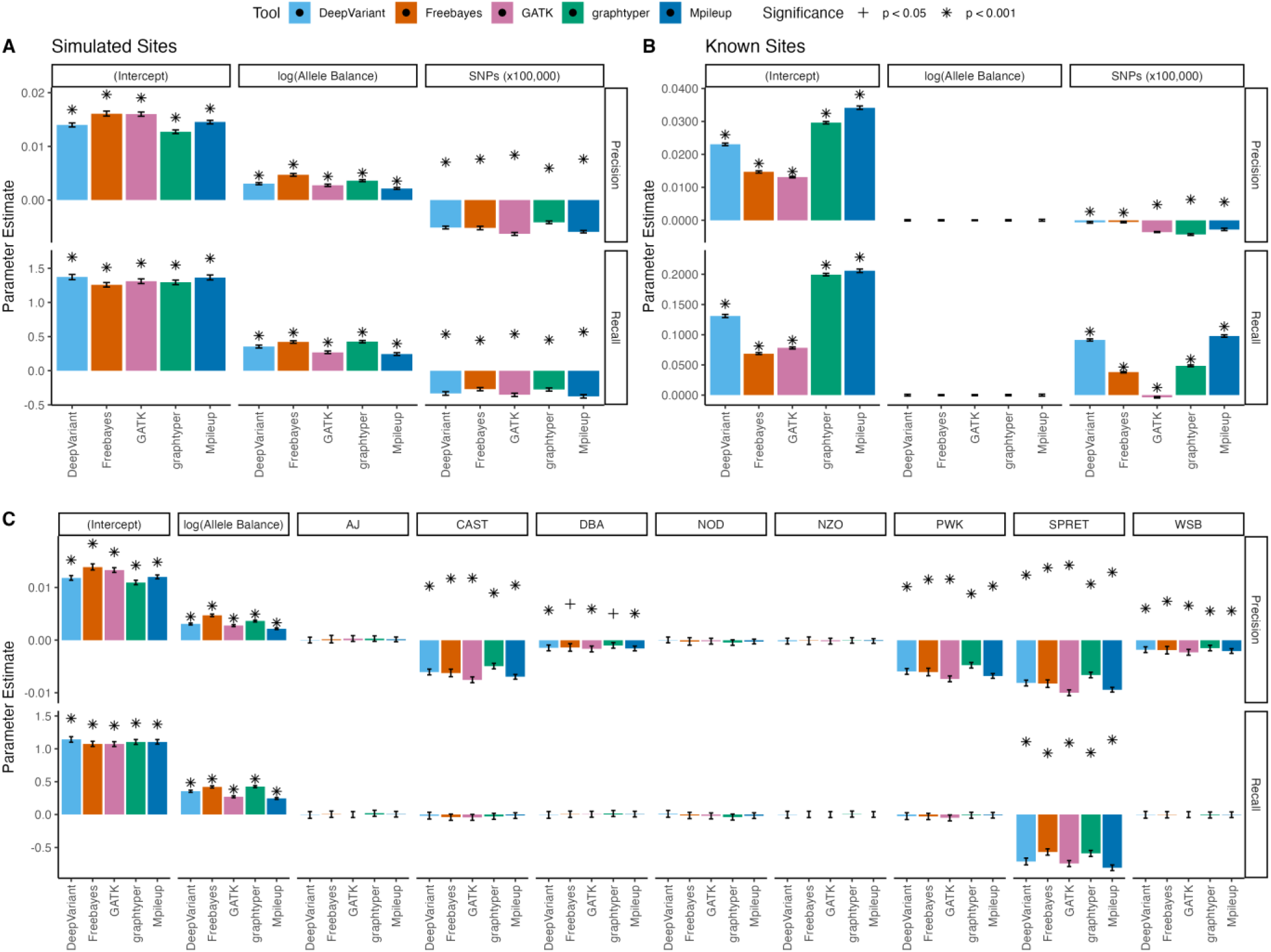
Comparison of coefficient estimates from linear models assessing the effect of log transformed allele balance and strain on precision and recall. (A and B) Coefficient estimates from linear models for each caller, with significance levels denoted. Strain effects were represented by the number of fixed differences relative to the chr1 GRCm39 reference sequence (coefficient for SNPs scaled by 100,000 for ease of visualization). Bars represent estimated effects, with error bars indicating ±2 standard errors across simulation replicates for (A) simulated sites and (B) known sites. Due to log-transformation of allele balance, model intercepts evaluate tool performance when allele balance is 1 and the focal strain is either identical to the reference genome (simulated sites, A) or identical to 129 (known sites, B). (C) Linear modeling results evaluating tool performance in the recovery of simulated sites, with strains treated as a factor and model coefficients calculated relative to 129, the strain with the lowest number of SNPs diverged from the reference.

Unsurprisingly, increasing SNP divergence relative to the reference genome reduces precision for all variant calling tools. We observe the largest negative impact with GATK (Figure 6A). Using 129 as a baseline (as the strain most closely related to the reference; Figure 5A), we find no significant difference in precision between 129 and A/J, NOD, or NZO, but model performance decreases for more divergent strains, with particularly steep losses for wild-derived strains CAST, PWK, WSB, and SPRET (Figure 6C). In contrast, the increasing number of SNPs improved the recall for known sites (Figure 6B). This observation could be due, at least in part, to a statistical effect: when both the number of predicted positives and the actual positives increase, the model has more opportunities to recover true positives, even without a substantive improvement in discrimination. Alternatively, this finding could indicate that a larger number of true positives increases the chance that some fall within regions of the genome where their detection is more accurate, effectively boosting true positive recovery. Below, we test this latter hypothesis by assessing model performance under various variant filtration conditions, including genomic positions and context.

#### 3.2.2 Evaluating Optimal Tool Combinations in Genetically Diverse Inbred Strains

As with B6, we used a decision tree to find the optimal combination of variant calling tools to produce a high-confidence variant call set for each of our genetically diverse inbred strains. Independent trees were constructed for each strain and AB combination and for both known and simulated sites. In all strains except SPRET and across all ABs, GATK had the highest importance values (Figure 7A). However, the relative importance of GATK compared to other callers did vary with AB. In the *Mus musculus domesticus* inbred strains (i.e., excluding CAST and PWK), the importance of DeepVariant was relatively higher at intermediate ABs (Figure 4A). In CAST and PWK, removing known sites had a greater importance on tree inference relative to other strains, in concordance with the higher number of fixed SNPs for those strains compared to the reference (Figure 7A, Figure 5B). Very few trees included Mpileup or graphtyper, and the importance of these callers was low in those trees that did retain these variant call outputs (Figure 7A).

**Figure 7.**
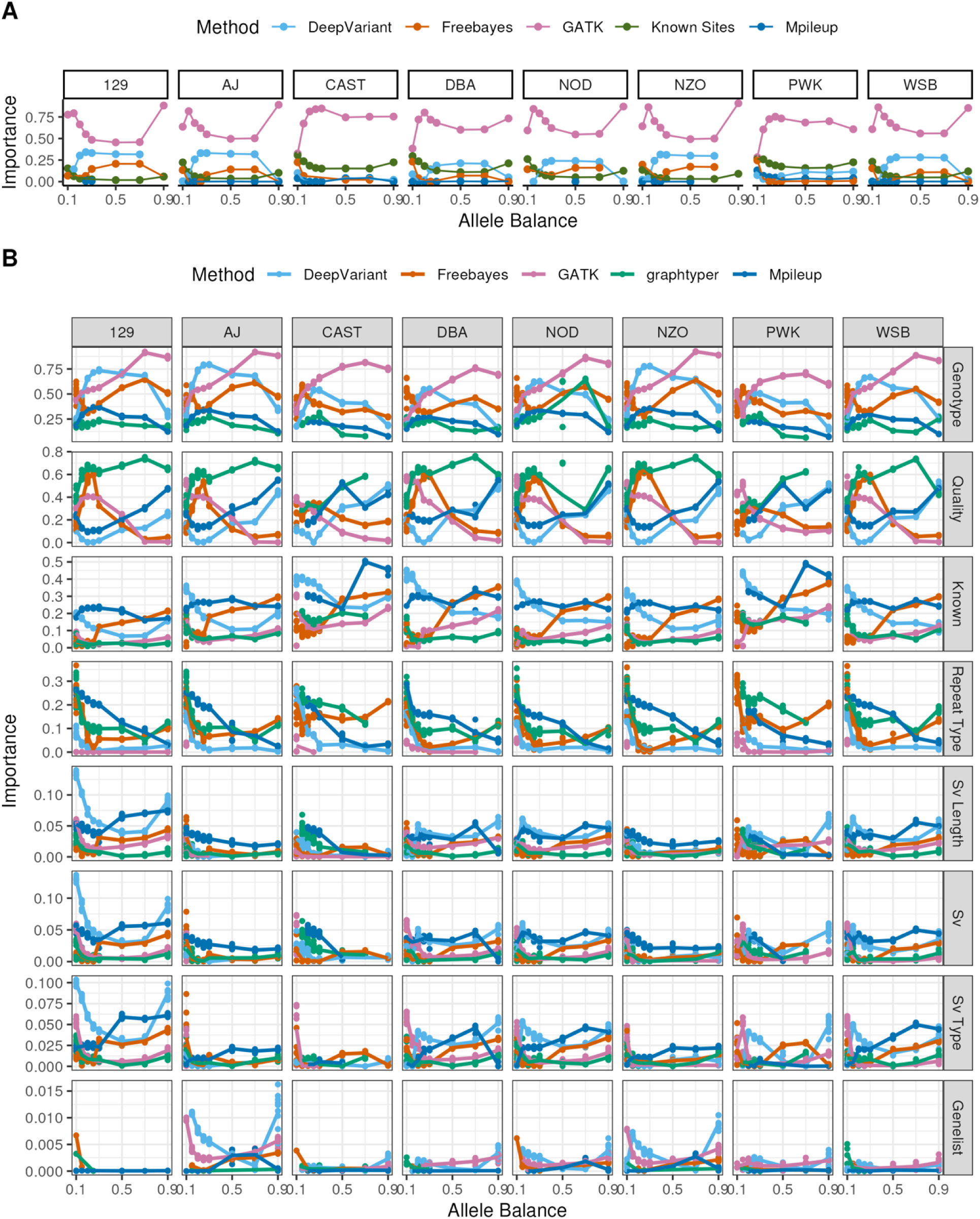
(A) Importance of variant calling tools in classification trees across different allele balances. Each line connects points representing the importance of a specific variant calling tool (indicated by line color) in classification trees predicting the true/false positive status of the variant. Note that the SPRET simulation has no points because no SPRET model had any supported splits based on included caller information. (B) Importance of filtration variables in classification trees across different allele balances for simulated sites across callers. Each line connects points representing the importance of a specific variable in classification trees predicting the true/false positive status of the variant. Panels are faceted in rows by the variable of interest, with one column for each strain. Points and lines are colored by the variant calling tool.

All SPRET trees had no splits, indicating that no combination of tools sufficiently alters the probability of recovering true positive variants enough to justify a split (see figure set S3). We suspect that this outcome reflects both the small number of simulated variants that were successfully lifted over from SPRET to the GRCm39 reference and the high number of false positive calls recovered in this strain across all tools (Figure 5).

### Site-Based Filtration Improves Precision Even in Highly Diverged Samples

To explore whether variant calling performance could be improved by applying genomic context or region-based filters, we built CART classification trees to predict whether called SNPs were part of the simulated SNP set for each tool, strain, and AB combination, using the following predictors: (i) genotype call state, (ii) variant quality, (iii) whether the site was a known site, (iv) whether the variant resides in an annotated repeat and if so, repeat type, (v) whether the variant lies a strain-specific structurally variable region, and if so, the length and type of SV, and (vi) overlap with a list of genes known to be rapidly evolving and structurally dynamic in mice. We find that all trees incorporated at least 2 filters, indicating that the performance of all tools can be improved with location-based filters (average number of rules across trees = 5.75, figure set S3). Trees for non-*domesticus* strains tended to feature more rules (average of 5.96 rules in PWK and 6.22 in CAST), and we report differences in the number of bifurcation rules across tools, ranging from 3.24 rules in GATK to 7.2 rules in Mpileup (figure set S3).

Across the assessed potential filtering criteria, the feature with the highest average importance was the genotype state output by the variant calling tool (e.g., heterozygous [0/1] or homozygous [0/1] for the alternate allele, as well as non-standard calls [1/0, 1|0, 0|1]), followed by variant quality and whether the location was a known site (Figure 7B). When filtering on genotype state, generally for known sites the trees recommended retaining only 1/1 sites, in alignment with our sequencing simulation workflow, while for simulated sites only 0/1, rather than homozygous or non-standard heterozygous call types were retained. Notably, the remaining features exhibited variability in their importance across replicates under a given set of simulation conditions, indicating sensitivity to the random location of simulated SNPs. Thus, it could be important to consider whether there are known mutational processes or biases influencing the genomic context of variants expected in a given study cohort, as this could inform the selection of optimal location-based filters. Generally, however, excluding SNPs in repeats had higher importance than any SV-related filter or eliminating SNPs in blacklisted genes (Figure 7B). This finding presumably reflects the larger number of bases annotated as repetitive DNA versus those resident within SVs or other blacklisted regions. The most common repeat family split across trees called for the removal of variants falling in low complexity, satellite, and simple repeats, as these regions are most enriched for false positives. The blacklisted gene list did not have high importance in our models, although few blacklisted genes are present on chr1. Future work exploring this feature in whole genomes is likely necessary to fully assess the utility of this filter (Figure 7B).

The application of these filters greatly increased the precision of the resulting call sets across all strains, eliminating a substantial proportion of the false positives and raising the precision of the naïve call sets (Figure 5D) to values above 0.5 for all strains, even at low ABs (Figure S5). However, this approach does reduce the recall considerably compared to the unfiltered call sets, particularly for extreme ABs. These findings emphasize the tradeoff between precision and recall under experimental conditions in variant calling. Post-filtration, the highest precision at low ABs is achieved using the GATK call sets. However, for ABs >0.5, the performance of DeepVariant and GATK are quite similar, with DeepVariant achieving higher precision in some conditions, and often with higher recall. Taken together, our findings indicate that straightforward filtration to exclude repetitive regions and strain-specific structural variation can improve the precision of variant calling without substantial reductions in recall for SNPs with intermediate ABs.

## Discussion

Variant calling must balance the detection of true variants while minimizing the recovery of false positive calls. Different variant calling tools address this goal through the implementation of distinct algorithms, statistics, and advised filtering approaches, but the relative performance of different callers in different settings is not fully understood. Here, we evaluate the overall performance and trade-offs among four popular variant calling tools using simulated data from inbred laboratory mouse strains. We find significant variability in the performance of our four tested callers, including differences in recall, precision, and false positive rates across different simulated ABs and coverage levels.

While depth of sequencing is known to impact SNP call rates (Kishikawa et al. 2019), we found that increasing depth had non-uniform effects on the performance of the 4 callers evaluated here. For example, at low coverage (10x), Mpileup had higher precision and recall than DeepVariant (Figure 3B), but at high coverage (40x), DeepVariant had higher precision than Mpileup. Similarly, higher ABs were associated with a higher probability of detection by all algorithms, but increasing AB had the largest effect on DeepVariant recall and the overall performance of Freebayes (Table 2).

Because the AB is expected to be 0.5 in heterozygous variants and 1 for variants fixed between inbred mouse strains, these findings are of deep importance to variant calling efforts in diverse laboratory mouse strains, including C57BL/6J. On the other hand, ABs may be much lower than 0.5 in studies of somatic variation, and caller performance at low ABs will have a large effect on the reliability of the resulting call sets and downstream biological inferences.

Our results suggest that the two haplotype-based variant calling approaches (GATK and Freebayes) generally out-perform alternative tools in precision but suffer a corresponding reduction in the recall. This may reflect the inherent properties of the callers themselves. For instance, the GATK HaplotypeCaller first flags potentially variable regions for local de novo reassembly. While this approach may improve accuracy, it may be overly stringent and miss a subset of true variants. While the other surveyed tools have lower precision, the application of straightforward, targeted filters can improve precision to within the ranges seen for Freebayes and GATK.

As has been suggested by other studies (Kim et al. 2014; Krøigård et al. 2016; Bian et al. 2018; Bergeron et al. 2022; Sendell-Price et al. 2023; Wooldridge et al. 2025), we find that an ensemble approach that relies on the integration of variants called by multiple variant calling tools produces superior results compared to any one tool on its own. However, our results do show that careful overlap approaches must be employed as some false positives are shared across multiple callers (Figure 4). Even in the reference strain, we recover some variants that are shared by three tools at low coverage (Figure S1,S2). At intermediate coverage levels in simulated data from B6, a combination of Mpileup and DeepVariant (Figure 4D) produces a high-quality call set with minimized false positives and the largest share of true positive SNPs. However, in the 40x coverage condition, most variants are recovered by all tools, and the lingering false positives detected by DeepVariant and Mpileup are likely counter-balanced by the recovery of unique true positive calls introduced by those tools (Figure S1,S2). Thus, while our work exposes a number of generalizable patterns, there is no single selection or combination of variant calling tools that will provide a one-size-fits-all solution for variant calling across all experimental and analytical applications in inbred house mice. For example, Freebayes output virtually no false positives in our simulated data (Figure 3A), contradicting a previous study reporting large numbers of false positive calls output by Freebayes in simulated human tumor datasets (Bian et al. 2018). Depending on the clinical or research question being pursued, Freebayes may be the most appropriate tool despite its absence in our optimal recommendations. Alternatively, if recall is more important than precision for a given analysis or question, exclusive use of DeepVariant or Mpileup may be the most beneficial approach.

Our work also underscores the importance of using data-driven filtering approaches. Genotype and the variant quality score were consistently the strongest predictor of variant validity, but genomic context also mattered. Notably, repetitive elements like low complexity, satellite, and simple repeats were enriched for false positives variant calls, highlighting the utility of straightforward repeat masking. While filtration clearly boosts precision, especially in low AB scenarios, it also reduces recall, reinforcing the inherent tradeoff in variant calling. Ultimately, our findings support a pragmatic approach: strategic use of genomic context, particularly repeats and known variant loci, can meaningfully improve variant calling precision without overly compromising recall, with varying magnitudes of importance depending on the types of variants being identified and strain genetic background.

Even after filtering, we recover a large number of persistent false positive SNP calls in divergent strains using Mpileup and Graphtyper. This finding likely points to deeper issues tied to cryptic strain-specific structural variation or alignment challenges that are not fully addressed by existing methods. In the case of graphtyper, some of these performance losses may be related to the underlying graph approach implemented by the software. Graph aligners try to construct paths in the alignment graph, but if samples carry large, novel variants not in the graph (e.g., insertions, rearrangements, or divergent haplotypes), reads may fail to align at all, align to the wrong part of the graph, or align equally well to multiple, incorrect paths. These outcomes could lead to missed variants, false positives, and overall poor call quality (Sibbesen et al. 2018; Hickey et al. 2020; Kaye and Wasserman 2021; Rautiainen et al. 2023). While we do not consider inversions here, we note that inverted regions may be especially important to investigate as regions enriched for poor-quality calls when using graph-based aligners in the future.

The utility of sequencing data depends on the availability and accuracy of tools to identify critical genomic features, including genetic variants. Through simulation, we show that selection of variant calling tools and filtration can have a meaningful impact on the accuracy of reported variants and that there are precision and recall trade-offs across callers. Because different clinical and scientific questions impose distinct tolerances for recall, precision, or false positives, different callers may be most appropriate in different experimental contexts. Through extensive simulation of sequencing data from the C57BL/6J inbred mouse strain and nine non-reference laboratory inbred models, we show that there are differences in SNP calling performance across tools, strains, and conditions, but that an optimized call set is obtained by using ensemble approaches and filtering based on genomic positions, with a particular eye towards repetitive regions. We provide a thorough release of the optimized workflow for each of the simulated contexts and strains we surveyed as supplementary material (Supp Reference). As the number of variant calling tools available for processing next-generation sequencing data continues to proliferate, careful benchmark evaluations are needed to understand their benefits and drawbacks and to set leading practices for the clinic and research environments. Further, understanding the conditions where existing tools underperform also allows for targeted development to fill existing software gaps.

---

**Supplemental datasheet 1: Accession numbers and info for strain references**

**Supplemental datasheet 2: All metrics B6**

**Supplemental datasheet 3: All metrics diverse strains**

**Supplemental datasheet 4: Full model results for non-reference strains**

**Code documents on figshare**

**Supplemental Figure S1:**
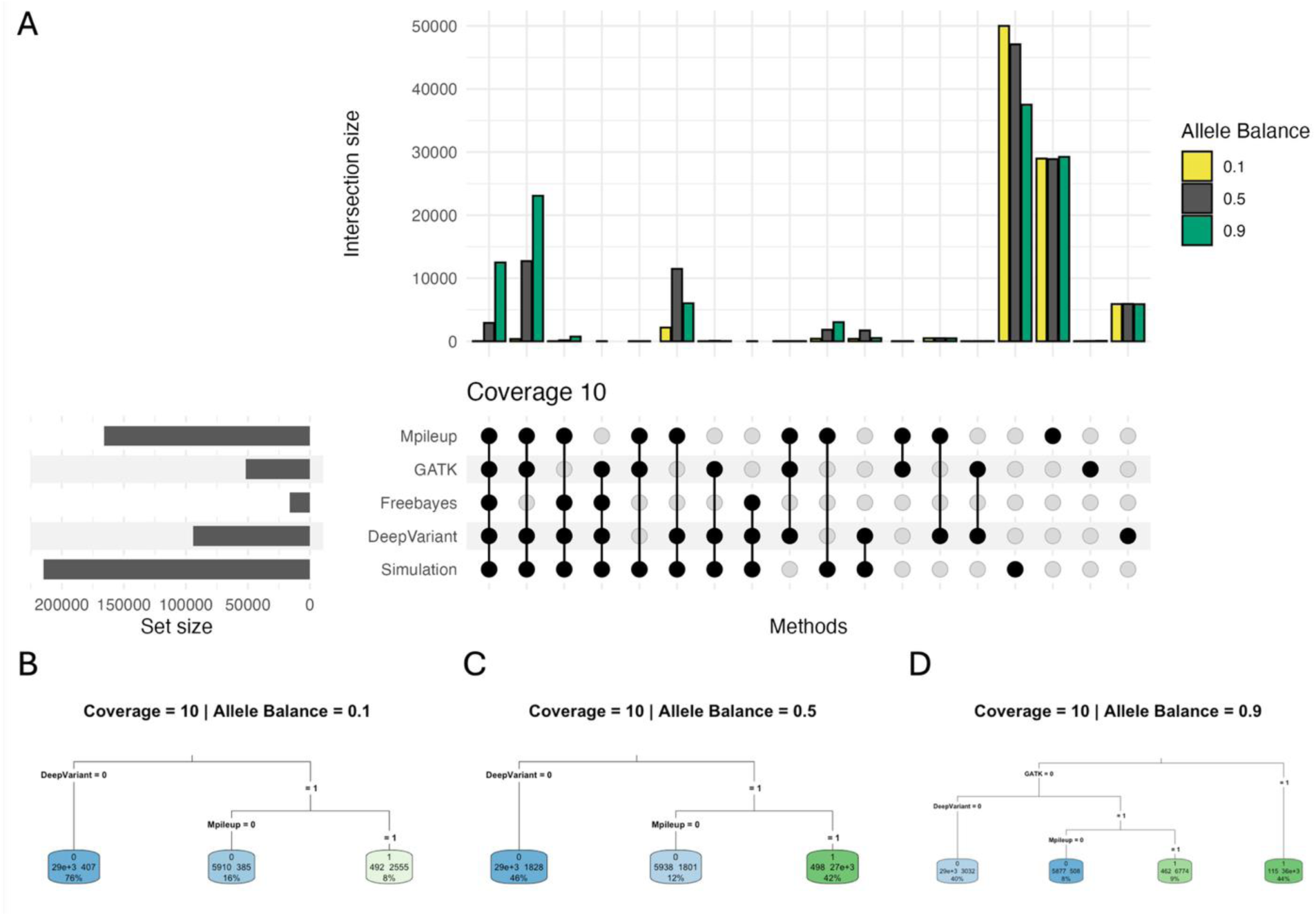
Comparisons of SNP call sets across tools. (A) Upset plot of variant call sets at 10x coverage and allele balances 0.1, 0.5, and 0.9. Bar graphs show the intersection sizes between each combination of tools, with each colored bar showing the size of that set at the given allele balance. Set sizes to the left of the upset chart panel indicate the number of variants recovered by that tool under the simulated allele balance condition. (B,C,D) Classification tree predicting variant status based on the optimal combination of variant calling tools under different allele balance conditions. Simulated variants are represented as “1”, and false positives are indicated by “0”. Splitting criteria are displayed on the connecting branches, indicating the decision rule applied to divide the data. Terminal (leaf) nodes show the final predicted value and sample size for each group of observations. Node color intensity represents the relative probability that an observation in that node belongs to the predicted class, with green indicating a higher probability (closer to 1) and blue indicating a lower probability (closer to 0). Darker colors correspond to more confident predictions, while lighter colors indicate greater uncertainty or class mixing within the node.

**Supplemental Figure S2:**
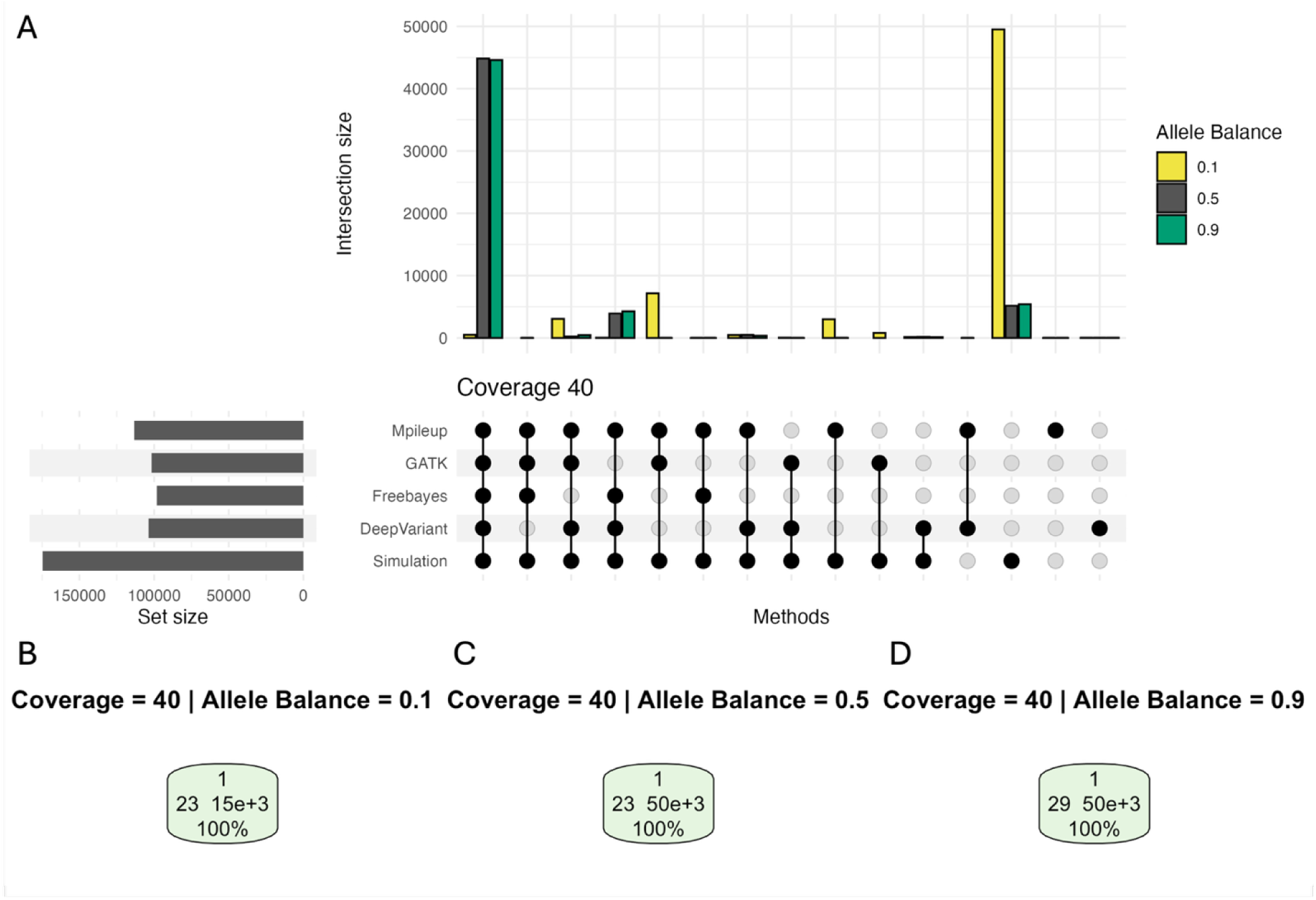
Comparisons of SNP call sets across tools. (A) Upset plot of variant call sets at 40x coverage and allele balances 0.1, 0.5, and 0.9. Bar graphs show the intersection sizes between each combination of tools, with each colored bar showing the size of that set at the given allele balance. Set sizes to the left of the upset chart panel indicate the number of variants recovered by that tool under the simulated allele balance condition. (B,C,D) Classification tree predicting variant status based on the optimal combination of variant calling tools under different allele balance conditions. Simulated variants are represented as “1”, and false positives are indicated by “0”. Splitting criteria are displayed on the connecting branches, indicating the decision rule applied to divide the data. Terminal (leaf) nodes show the final predicted value and sample size for each group of observations. Node color intensity represents the relative probability that an observation in that node belongs to the predicted class, with green indicating a higher probability (closer to 1) and blue indicating a lower probability (closer to 0). Darker colors correspond to more confident predictions, while lighter colors indicate greater uncertainty or class mixing within the node.

**Supplemental Figure Set 3: All trees B6, all trees diverse strains.**

**Supplemental Figure S4.**
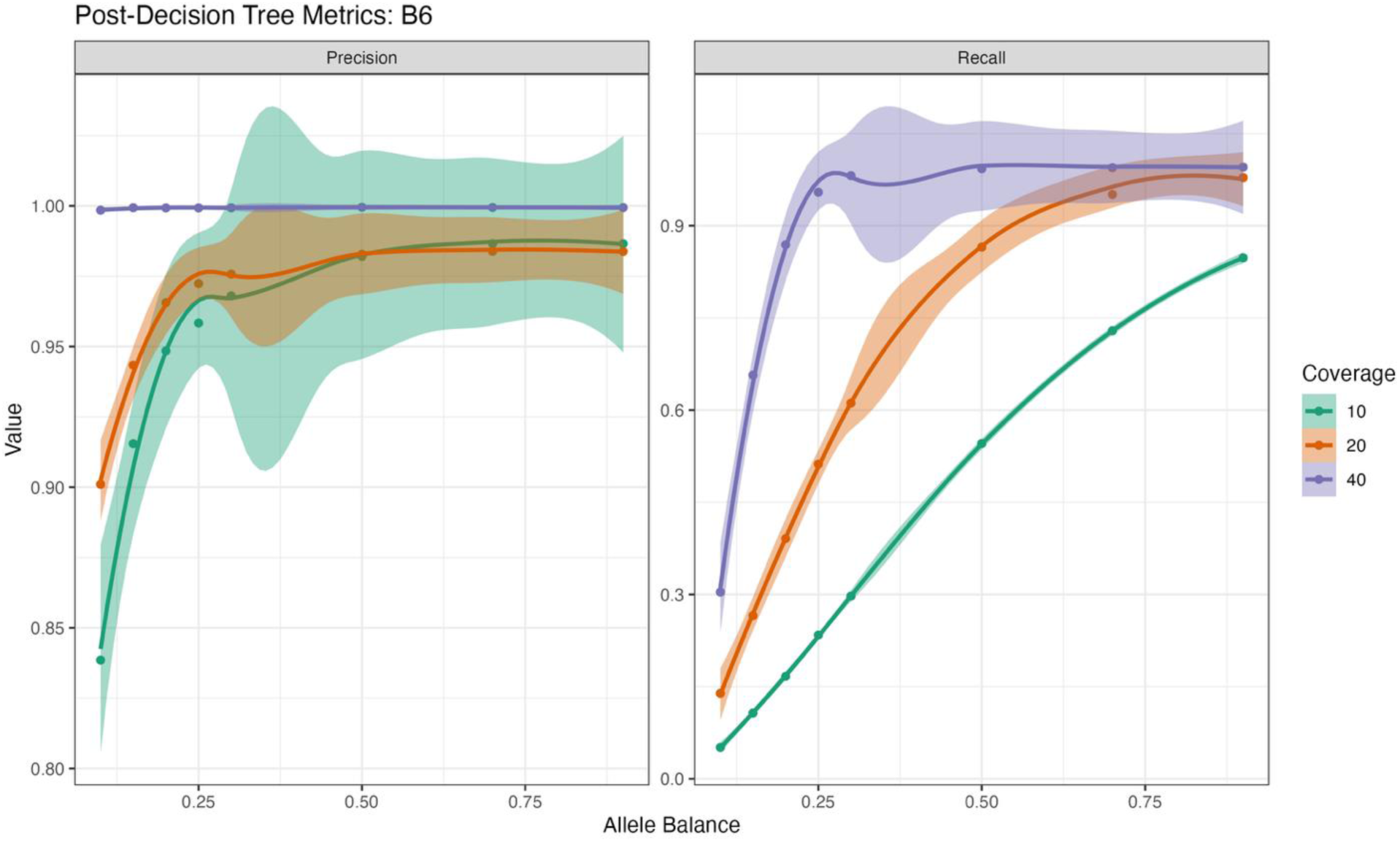
Post optimal filtration. variant calling tool performance comparison across all allele balance and coverage values for Precision and Recall.

**Supplemental Figure S5.**
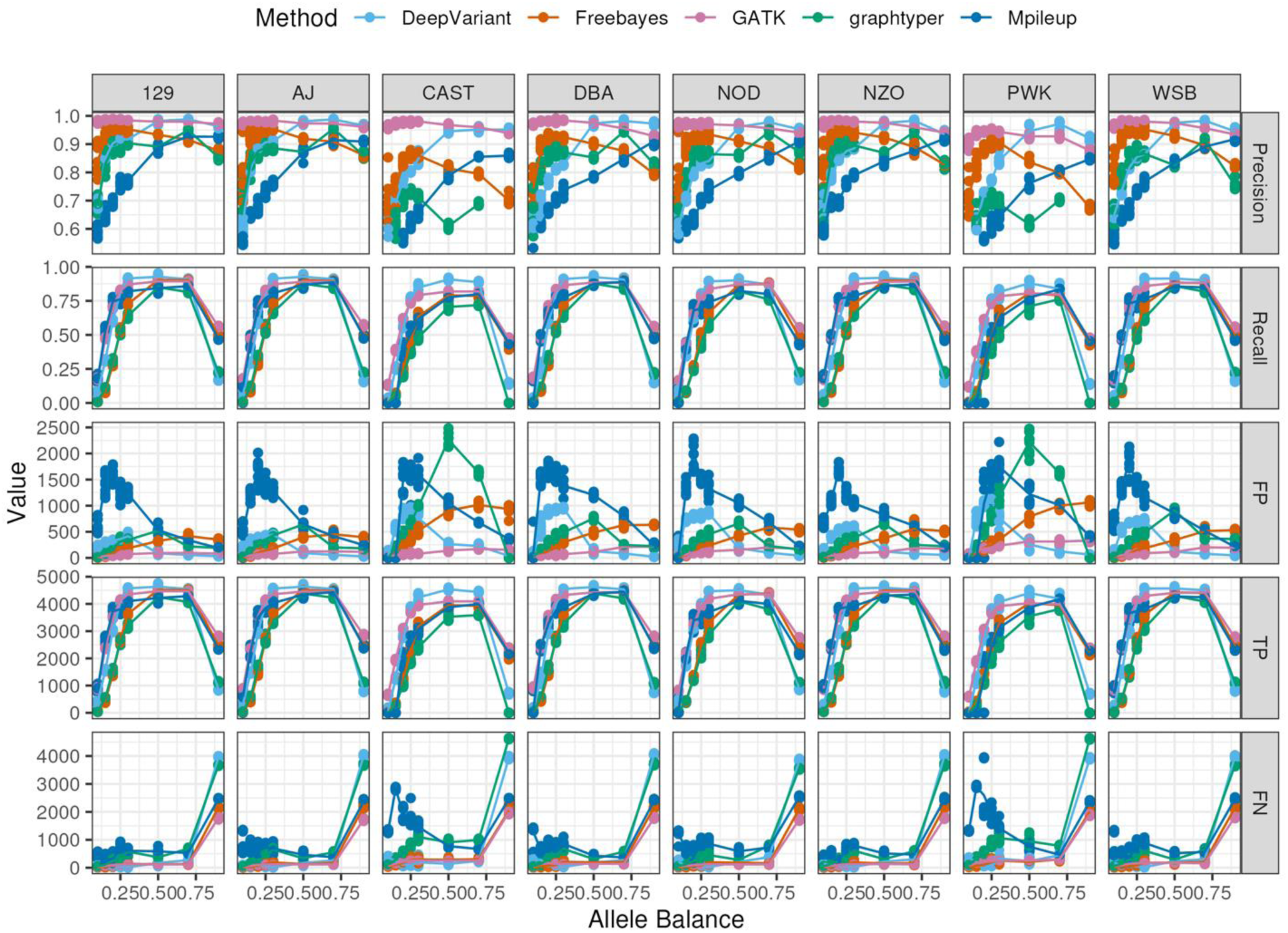
Post optimal filtration. variant calling tool performance comparison across all allele balance and strains by Precision, Recall, FP (the number of False Positive calls remaining in the optimal dataset), TP (the number of True Positive calls remaining after filtration), and FN (variants initially called correctly, but subsequently eliminated by the filtration approach).

## Acknowledgments

The authors gratefully acknowledge the Research IT department at The Jackson Laboratory for their support and assistance in this work and the maintenance of the high-performance computing resources used for conducting the research reported in this paper. The authors thank the Dumont Lab members, particularly Lydia Wooldridge, for their critical review and helpful discussions, which contributed to refining this manuscript. We also acknowledge the support of the Tufts Writing Center, and Christine Smith, for providing editing support.

## Author Contributions

AG, LBB, and BLD initiated and conceptualized the research project. LBB led the investigation, including simulations and variant calling, supported by AG and BLD. AR contributed to the investigation and formal analysis of the non-B6 strains. AG led the formal analysis and visualization, supported by LBB and BLD. BLD provided supervision and resources throughout the project. AG wrote the original manuscript draft, and LBB and BLD provided review and edits. All authors have read and approved the final manuscript.

## Data Availability

The datasets and analysis code supporting the findings of this study are available on Figshare at www.doi.org/10.6084/m9.figshare.29089805 [ embargoed, private link: https://figshare.com/s/27f2e65c473f52c2e311 ]. This repository includes all processed data, analysis scripts, and supplementary materials necessary to reproduce the results reported in the manuscript. The references used in generating the non-B6 sequencing are available under the following accession numbers: GCA_001624445.1, GCA_001624835.1, GCA_001624775.1, GCA_001624505.1, GCA_001624745.1, GCA_001624185.1, GCA_001624215.1, GCA_001624675.1, and GCA_921997135.2.

